# An elite broadly neutralizing antibody protects SARS-CoV-2 Omicron variant challenge

**DOI:** 10.1101/2022.01.05.475037

**Authors:** Biao Zhou, Runhong Zhou, Jasper Fuk-Woo Chan, Mengxiao Luo, Qiaoli Peng, Shuofeng Yuan, Bobo Wing-Yee Mok, Bohao Chen, Pui Wang, Vincent Kwok-Man Poon, Hin Chu, Chris Chung-Sing Chan, Jessica Oi-Ling Tsang, Chris Chun-Yiu Chan, Ka-Kit Au, Hiu-On Man, Lu Lu, Kelvin Kai-Wang To, Honglin Chen, Kwok-Yung Yuen, Zhiwei Chen

## Abstract

The strikingly high transmissibility and antibody evasion of SARS-CoV-2 Omicron variant have posted great challenges on the efficacy of current vaccines and antibody immunotherapy.Here, we screened 34 BNT162b2-vaccinees and cloned a public broadly neutralizing antibody (bNAb) ZCB11 from an elite vaccinee. ZCB11 neutralized all authentic SARS-CoV-2 variants of concern (VOCs), including Omicron and OmicronR346K with potent IC50 concentrations of 36.8 and 11.7 ng/mL, respectively. Functional analysis demonstrated that ZCB11 targeted viral receptor-binding domain (RBD) and competed strongly with ZB8, a known RBD-specific class II NAb. Pseudovirus-based mapping of 57 naturally occurred single mutations or deletions revealed that only S371L resulted in 11-fold neutralization resistance, but this phenotype was not observed in the Omicron variant. Furthermore,prophylactic ZCB11 administration protected lung infection against both the circulating pandemic Delta and Omicron variants in golden Syrian hamsters. These results demonstrated that vaccine-induced ZCB11 is a promising bNAb for immunotherapy against pandemic SARS-CoV-2 VOCs.

## INTRODUCTION

After two years of the COVID-19 pandemic, the highly transmissible SARS-CoV-2 and its variant of concerns (VOCs) have resulted in more than 279 million infections with 5.4 million deaths globally by December 26, 2021 (https://coronavirus.jhu.edu/map.html). During this period, various types of COVID-19 vaccines have been quickly developed to control the pandemic with over 8.9 billion doses administered in many countries. Although the extensive implementation of vaccination has significantly reduced the rates of hospitalization, severity and death ^1-5^, current vaccines do not confer complete or durable prevention of upper airway transmission of SARS-CoV-2. The numbers of vaccine-breakthrough infections and re-infections, therefore, have been continuously increasing ^6-8^. The pandemic situation has been complicated by repeated emergence of new VOCs, including Alpha (B.1.1.7), Beta (B.1.351), Gamma (P.1), Delta (B.1.617.2) and Omicron (B.1.1.529) ^9,10^, and waning of vaccine-induced immune responses, together with relaxed preventive masking and social distancing ^11-13^.

After the World Health Organization (WHO) designated the Omicron as a VOC on November 26, 2021, this variant has been quickly found in over 110 countries and is replacing the Delta VOC within a month, becoming the dominant VOC in many places in the South Africa, European countries, and the United States ^14,15^. According to the GASAID database, for example, the relative variant genome frequency of the current circulating Delta VOC has declined from 89% to 19.6% while the Omicron VOC has increased from 0% to 67.4% in African countries during the period from October 4, 2021, to December 26, 2021. The rapid global spread of the Omicron VOC has been associated with vaccine-breakthrough infections and re-infections ^16,17^. Moreover, like previous findings that the Beta VOC compromised vaccine-induced neutralizing antibody (NAb) ^12,18,19^, the Omicron VOC has resulted in even worse NAb evasion due to more than 30 alarming mutations in SARS-CoV-2 spike glycoprotein ^20-23^. Considering that current NAb combination for clinical immunotherapy showed significantly reduced activities ^21,24^, we sought to search for vaccine-induced broadly neutralizing antibody (bNAbs) among elite vaccinees.

## RESULTS

### Identification of an elite vaccinee who developed bNAbs

To isolate potent bNAbs against currently circulating SASR-CoV-2 VOCs, we searched for elite vaccinees, who had developed potent bNAbs among a Hong Kong cohort of 34 vaccinees, around average 30.7 days (range, 7-47 days) after their second dose of the BNT162b2 vaccination (BioNTech-Pfizer) (Supplementary Table 1) ^13^. 100% subjects developed NAbs against the pseudotyped SARS-CoV-2 wildtype (WT, D614G) (Fig. 1a, top left). To seek for vaccinees with bNAb, we then tested the full panel of pseudotyped SARS-CoV-2 VOCs including Alpha (B.1.1.7), Beta (B.1.351), Gamma (P.1), Delta (B.1.617.2) and Omicron (B.1.1.529). Only two study subjects (2/34), BNT162b2-26 and BNT162b2-55, were considered as elite vaccinees who harbored bNAbs with IC_90_ or IC_50_ values higher than the mean titers of all VOCs tested in the cohort. BNT162b2-26 displayed significantly high bNAbs titers against the Beta and Delta variants (Fig. 1a, mid left and bottom left, Supplementary Table 2), the known most resistant VOC and the dominant VOC, respectively, before the Omicron variant ^25,26^. After measuring binding antibodies to spike protein (Fig. 1b), we calculated the neutralizing potency index as previously described ^27^. We found that Omicron resulted in the highest reduction of the mean neutralizing potency index as compared with other VOCs (Fig. 1c). BNT162b2-26, however, displayed neutralizing potency index scores consistently higher than the mean ones against all VOCs tested. We, therefore, chose this elite vaccinee for subsequent search of bNAb.

**Fig. 1.**
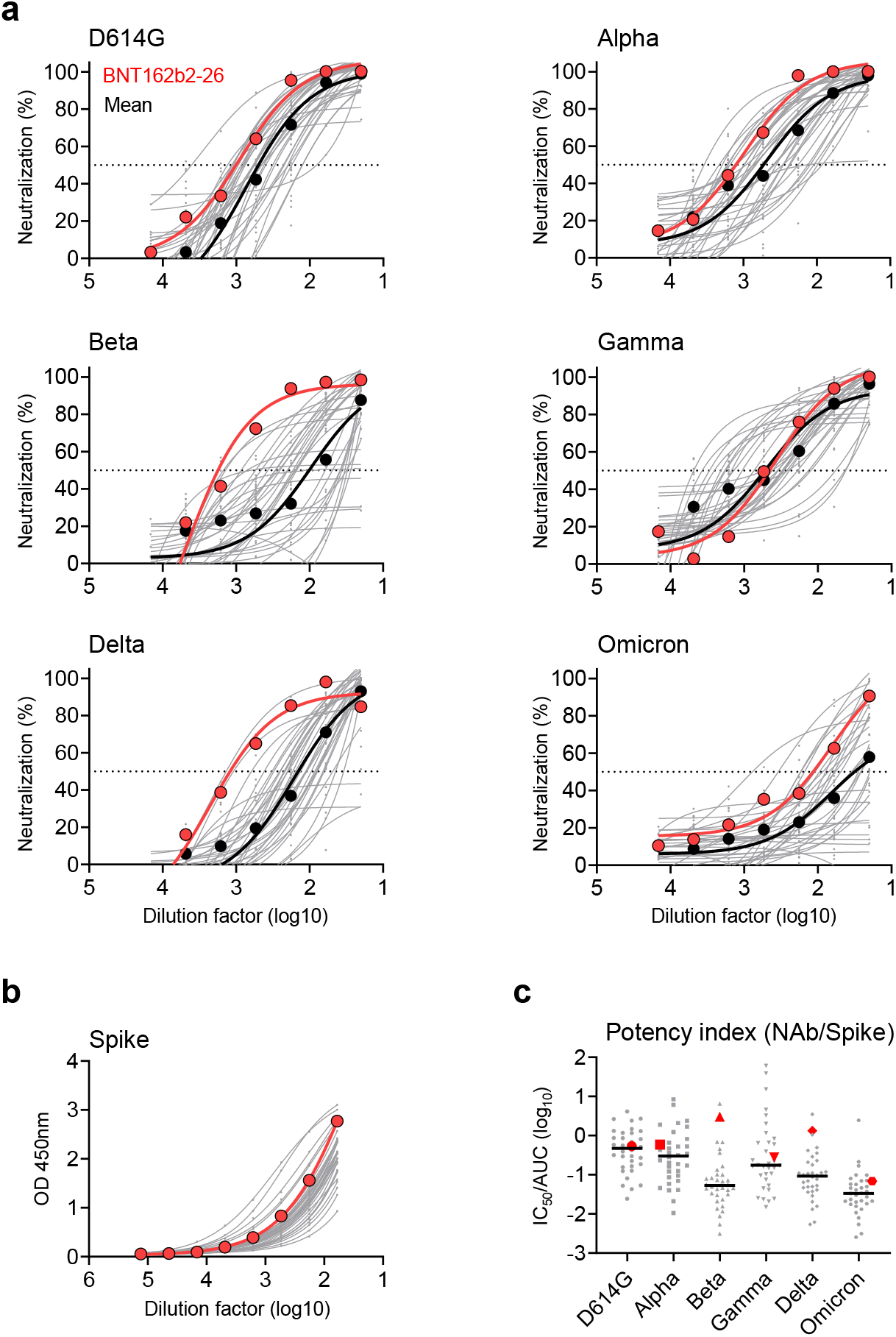
Identification of an elite vaccinee who developed bNAbs. Plasma samples derived from 34 BNT162b2-vaccinees were tested at average 30.7 days (range 7-47 days) after second vaccination (BioNTech-Pfizer). (**a**) Serially diluted plasma samples were subjected to neutralization assay against the pseudotyped SARS-CoV-2 WT and five variants of concern. The neutralizing curve of the elite BNT162b2-26 vaccinee (red) was compared with the mean curve of all vaccinees tested (dark black). (**b**) Binding activity of spike-specific plasma IgG was determined by ELISA at serial dilutions. The binding curve of the elite BNT162b2-26 vaccinee was presented as red. (**c**) The neutralization antibody potency index was defined by the ratio of IC_50_/AUC of anti-spike IgG in BNT162b2-vaccinees. Neutralizing IC_50_ values represented plasma dilution required to achieve 50% virus neutralization. The area under the curve (AUC) represented the total peak area was calculated from ELISA OD values. Each symbol represented an individual vaccinee with a line indicating the median of each group. The elite BNT162b2-26 vaccinee was presented as red dots.

### Isolation of NAbs against SARS-CoV-2 from the elite vaccinee

With vaccinee informed consent, we obtained another blood sample donated by BNT162b2-26 at day 130 after his second vaccination. Fresh PBMCs from BNT162b2-26 were stained for antigen-specific memory B cells (CD19, CD27, IgG) using the 6xHis-tagged SARS-CoV-2 WT spike as the bait as previously described ^28^. Spike-specific memory B cells were found in BNT162b2-26 but not in the healthy donor (HD) control (Supplementary Fig. 1) and were sorted into each well with a single B cell for antibody gene amplification. After antibody gene sequencing, we recovered 14-paired heavy chain and light chain for antibody IgG1 engineering. Seven of these 14 paired antibodies including ZCB3, ZCB8, ZCB9, ZCB11, ZCC10, ZCD3, ZCD4 in antibody expression supernatants showed positive responses to WT spike by ELISA 48 hours post transient transfection (Supplementary Fig. 2a). Five of these seven spike-reactive antibodies including ZCB3, ZCD4, ZCB11, ZCC10 and ZCD3 targeted spike S1 subunit (Supplementary Fig. 2b), whereas ZCB8 and ZCB9 were S2-specific (Supplementary Fig. 2c). Moreover, among these five S1-reactive antibodies, only ZCD4 was not specific to RBD (Supplementary Fig. 2d) and none of them interacted with NTD (Supplementary Fig. 2e, Supplementary Table 3). Eventually, only four RBD-specific ZCB3, ZCB11, ZCC10 and ZCD3 showed neutralizing activities against WT by the pseudovirus neutralization assay (Supplementary Fig. 2f). These results demonstrated that RBD-specific NAbs were primarily obtained from memory B cells of BNT162b2-26 at 130 days after his second vaccination.

Notably, besides the previously published control ZB8 ^28^, ZCB11 had the strongest binding capability to both RBD and Spike with the same EC_50_ values of 20 ng/ml by ELISA (Fig. 2a, Supplementary Table 5). Moreover, the binding dynamics of ZCB11 to SARS-CoV-2 RBD was determined using the surface plasmon resonance (SPR). We found that ZCB11 exhibited the fast-on/slow-off kinetics with an equilibrium dissociation constant (KD) value of 5.75×10^−11^ M, suggesting an RBD-specific high-binding affinity (Supplementary Fig. 2g and Supplementary Table 6). In subsequent quantitative neutralization analysis against WT, we found that two of these four NAbs, ZCB3 and ZCB11, showed high neutralization potency with IC_50_ values below 100 ng/mL (Fig. 2b top left, Supplementary Table 7). Sequence analysis revealed that ZCB3, ZCC10 and ZCD3 utilized IGHV3-53/3-66 heavy chain, whereas their paired light chains had distinct IGKV1-9, IGKV3-20 and IGKV1-27, respectively (Supplementary Table 3). In contrast, ZCB11 utilized different IGHV1-58 heavy chain and IGKV3-20 light chain. Our four new NAbs were all considered as public antibodies characterized by a IGHV3-53/3-66 heavy chain with 10-12 residues in the CDR3 region or a IGHV1-58 heavy chain with 15-17 residues in CDR3 region as previous reported by others ^12,29,30^. These results demonstrated BNT162b2-26 developed mainly public NAbs after two doses of vaccination.

**Fig. 2.**
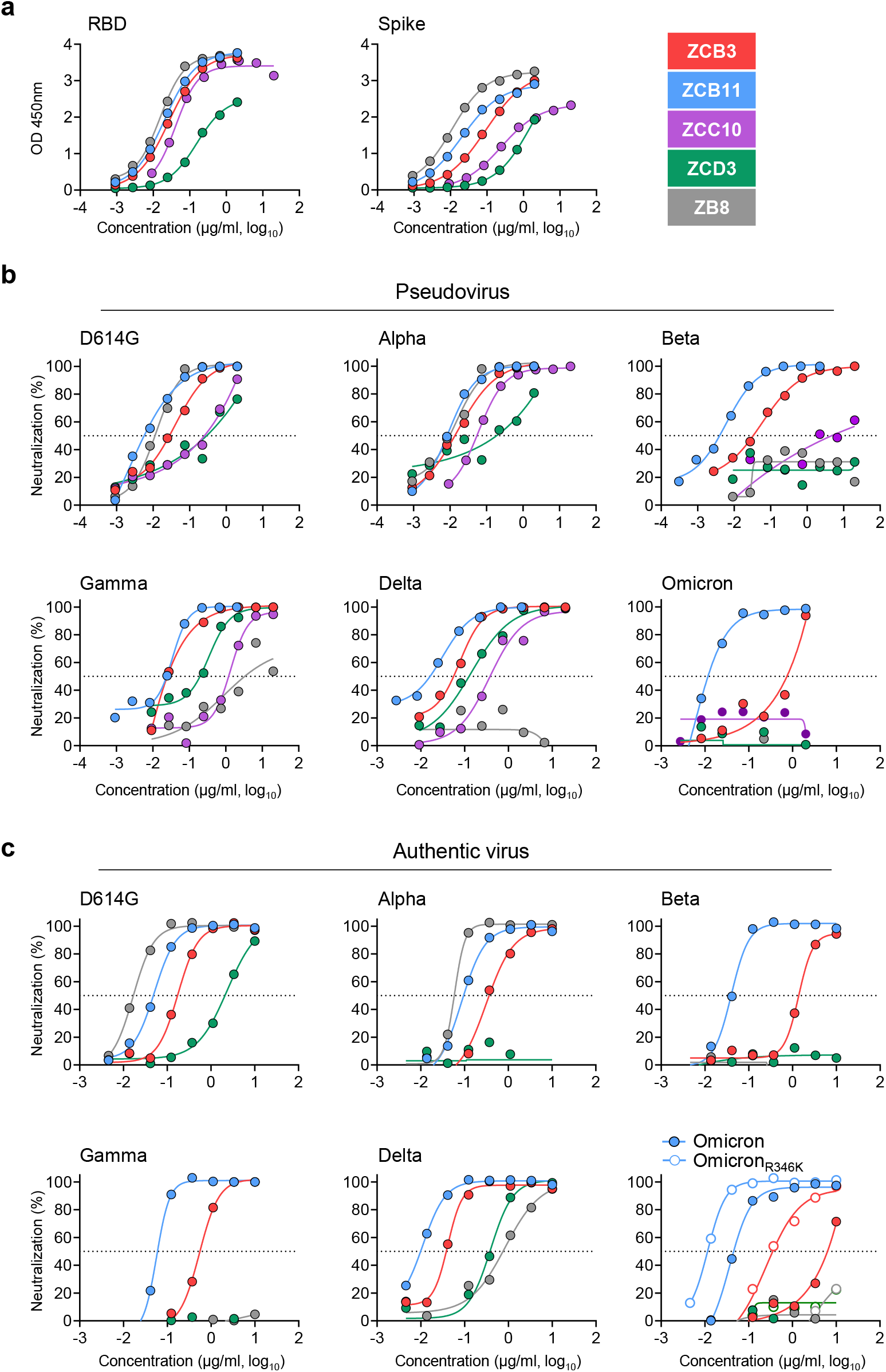
Comparison of bNAbs isolated from the elite vaccinee. (**a**) RBD-and spike-specific binding activities of 4 newly cloned NAbs including ZCB3, ZCB11, ZCC10 and ZCD3 were determined by ELISA at serial dilutions. A known NAb ZB8 was included as a control. (**b**) Neutralizing activities of ZCB3, ZCB11, ZCC10 and ZCD3 were determined against six pseudotyped SARS-CoV-2 variants of concern including D614G (WT), Alpha, Beta, Gamma, Delta and Omicron as compared with the control NAb ZB8. (**c**) Neutralizing activities of ZCB3, ZCB11, ZCC10 and ZCD3 were determined against the same six but authentic SARS-CoV-2 variants of concern as compared with the control NAb ZB8. The color coding was consistently used in **a-c**. Notably, the authentic SARS-CoV-2 Omicron_R346K_ was also tested (bottom right in **c** with empty dots). The dashed line in each graph indicated 50% neutralization.

### Antibody neutralization of SARS-CoV-2 VOCs

To understand the breadth of these four newly cloned public RBD-specific NAbs, we performed SARS-CoV-2 neutralization assays using both pseudoviruses and authentic VOC isolates, including Alpha, Beta, Gamma, Delta and Omicron variants (Fig. 2b). ZB8, a known RBD-specific class II NAb, was included as a positive control. Testing pseudoviruses in 293T-ACE2 cells, we found that ZCB11 was the best bNAb that neutralized all VOCs potently, including the most alarming Omicron variant ^21^, with IC_50_ values of around 30 ng/mL for Gamma and Delta variants and 6 ng/mL for Alpha, Beta and Omicron variants (Fig. 2b, Supplementary Table 7). ZCB3 was the second best bNAb and neutralized Alpha, Beta, Gamma and Delta variants potently, but not the Omicron variant. ZCC10 and ZCD3 neutralized Alpha, Gamma and Delta variants at relative low potency, but lost neutralization totally against Beta and Omicron variants. Importantly, testing authentic VOC viruses in Vero-E6-TMPRSS2 cells, we consistently found that ZCB11 was the most potent bNAb, followed by ZCB3 (Fig. 2c). The IC_50_ values of ZCB11 for neutralizing Alpha, Beta, Gamma, Delta, Omicron and Omicron_R346K_ variants were 85.1, 39.9, 56.9, 11.2, 36.8 and 11.7 ng/mL, respectively, which were comparable to the IC_50_ value of 51 ng/ml for neutralizing the WT (Supplementary Table 7). ZCB3 was about 10-fold less potent than ZCB11 for neutralizing Beta and Omicron variants. Notably, the potency of ZCB11 in the pseudovirus assay was higher than that in the authentic virus assay, which was probably related to different target cells used. ZB8 showed unmeasurable and weak neutralization against Delta pseudovirus and Delta authentic virus, respectively. Conversely, ZCC10 and ZCD3 showed weak and unmeasurable neutralization against Gamma pseudovirus and Gamma authentic virus, respectively. These results demonstrated that ZCB11 functioned as an elite bNAb potently neutralized all circulating SARS-CoV-2 VOCs *in vitro*. Notably, although BNT162b2-26 developed mainly public NAbs, ZCB11 was unlikely dominantly elicited due to the reduced titer against Omicron as compared with WT (Fig. 1).

### Naturally occurred mutations or deletions conferring antibody resistance

Since Omicron variant escaped from NTD-specific NAbs and majority of known RBD-specific NAbs in the class I, class II, class III and class IV groups ^20,21,31^, we sought to determine possible mutations or deletions responsible for antibody resistance for ZCB3 and ZCB11 as compared with the control ZB8. We first constructed and tested a large panel of pseudoviruses carrying individual mutations or deletions found in Omicron variant as compared with those previously found in Alpha, Beta, Gamma and Delta variants (Fig. 3a). For ZB8, we consistently found that the E484 is essential for its neutralization activity. E484K in Beta, E484Q in Delta and E484A in Omicron were responsible for the significant ZB8 resistance, followed Q493R for about 10-fold resistance. For ZCB3, none of single mutations or deletions tested conferred resistance for equal to or more than 10-fold. Only and Q493R in Omicron reduced neutralization potency of around 3.5-fold. For ZCB11, only S371L in Omicron showed 11.2-fold resistance (Fig. 3a). Moreover, Q493R, Y505H, T547K and Q954H in Omicron exhibited over 6-fold resistance. Unexpectedly, when all these and other mutations combined in Omicron, they did not confer significant resistance at all. Subsequently, we performed antibody competition by Surface SPR. Although they engaged different clonotype and antibody resistant profiles, ZCB11 exhibited as a strong competitor for WT RBD binding against either ZCB3 or ZB8, respectively (Fig. 3b and Supplementary Fig. 3), suggesting overlapped antibody binding epitopes in RBD between them. To further predict the binding mode of ZCB11, we searched structural database for RBD-specific NAbs with similar B cell clonotype. Interestingly, the patient-derived S2E12 Nab, which used the same IGHV1-58 heavy chain and IGKV3-20 light chain ^32^, shared the high amino acid identity of 82.2% in heavy chain variable regions with ZCB11. A model of ZCB11 variable regions was generated based on the protein sequence by the SWISS-MODEL using the crystal structure of S2E12 Fab fragment (Research Collaboratory for Structural Bioinformatics [RCSB] PDB code 7K3Q) as the template. The superimposed ZCB11 and S2E12 variable regions (Fig. 3c) showed that the secondary elements and most of loops are relatively conserved, except for the HCDR1 and KCDR3 which contained a single amino acid insertion and deletion, respectively. It is possible that ZCB11 also recognized the convex receptor binding motif (RBM) like S2E12. S477N, Q493R and Y505H mutations that conferred partial ZCB11 resistance in the pseudovirus assay were close to the binding interface between S2E12 and RBM (Fig. 3d).

**Fig. 3.**
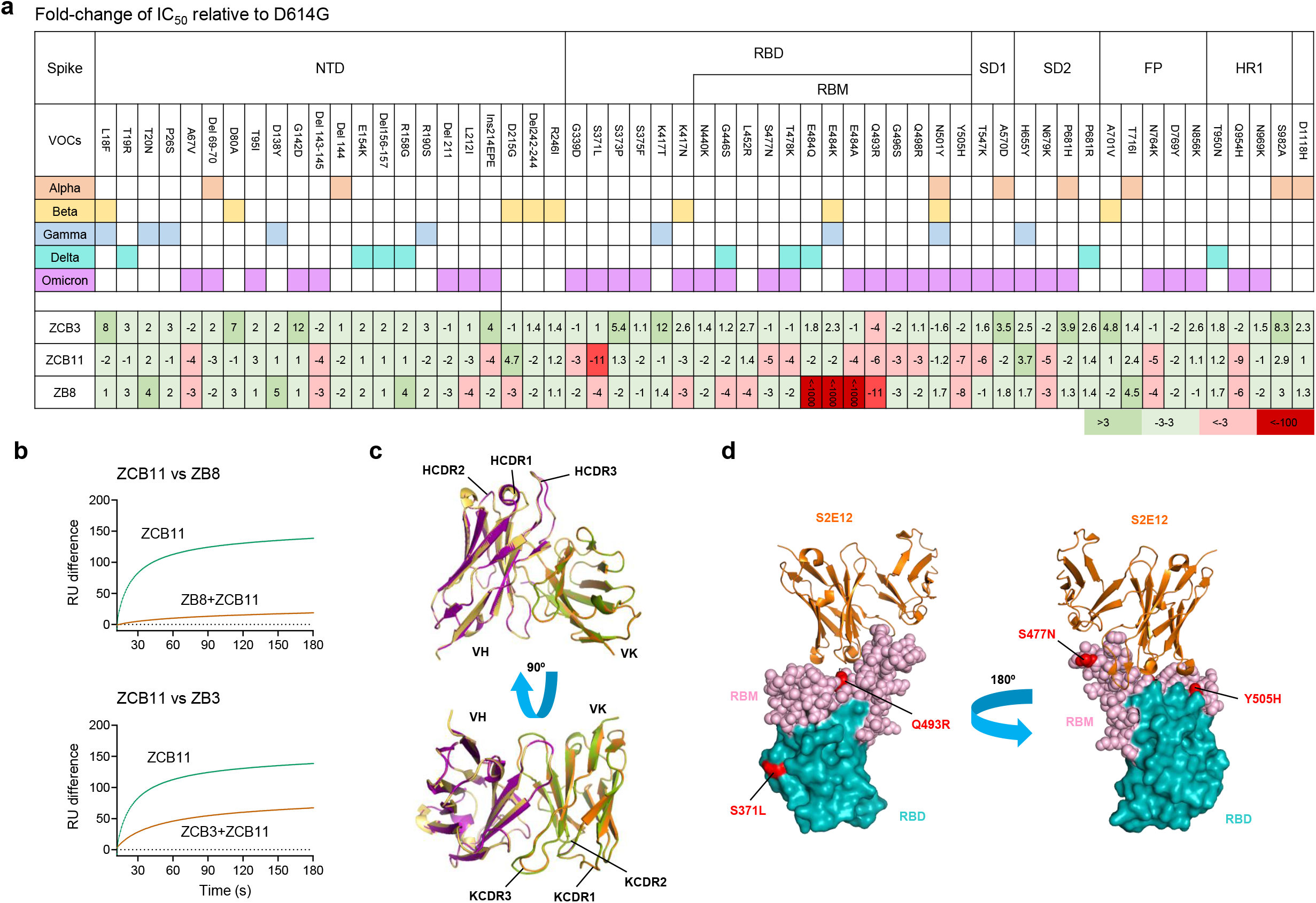
Naturally occurring mutations or deletions conferring antibody resistance in VOCs. (**a**) Fold change of IC_50_ values relative to WT was determined by pseudoviruses carrying individual mutations or deletion against bNAbs ZCB3 and ZCB11 as compared with ZB8. (**b**) Antibody competition by SPR between ZCB11 and ZC8 (top) as well as between ZCB11 and ZCB3 (bottom). (**c**) Structural alignment between S2E12 and ZCB11 variable regions. The structure of the ZCB11 variable region predicted by the SWISS-MODEL is superimposed into the structure of S2E12 (PDB: 7K3Q). Cartoon representation of ZCB11 variable region of heavy chain (VH) is shown in purple and the variable region of light chain (VK) in orange. The S2E12 VH and VK are shown in yellow and green, respectively. The CDRs of VH and VK are labelled. (**d**) The structure of RBD in complex with the S2E12 variable region (from PDB 7K45). RBD is shown in cyan with receptor binding motif (RBM) highlighted in light pink and the amino acids whose substitution confers resistance to ZCB11 in (**a**) are highlighted in red.

### *In vivo* efficacy of ZCB11 against SARS-CoV-2 Delta and Omicron variants

To determine the *in vivo* potency of ZCB11 against the dominant circulating VOCs, we conducted viral challenge experiments using the golden Syrian hamster COVID-19 model as compared with ZB8 ^33^. Since ZB8 conferred nearly complete lung protection against SARS-CoV-2 WT intranasal challenge at 4.5 mg/kg as we previously described ^34^, we tested it in parallel with ZCB11 using the same dose according to our standard experimental procedure (Fig. 4a). One day prior vial challenge, three groups of hamsters (n=8) received the intraperitoneal injection of ZCB11, ZB8 and PBS, respectively. Twenty-four hours later, half of the animals (n=4) in each group were separated into subgroups and were challenged intranasally with 10^5^ PFU of SARS-CoV-2 Delta variant and Omicron variant, respectively. Animal body weight changes were measured daily until day 4 when all animals were sacrificed for endpoint analysis. For three subgroups challenged with the SASR-CoV-2 Delta variant, we found that the infection caused around 10% body weight loss overtime in the PBS and ZB8 pre-treatment groups. In contrast, transient and less than 4% body weight decrease was observed for the ZCB11-treated hamsters (Fig. 4b). Moreover, relatively lower sub-genomic viral loads (Fig. 4c) and unmeasurable numbers of live infectious viruses (six orders of magnitude drop) (Fig. 4d) were achieved by ZCB11 than by ZB8. For hamsters challenged with the SASR-CoV-2 Omicron variant, no significant body weight loss was found in all three subgroups, indicating relatively weaker pathogenicity caused by Omicron than by Delta (Fig. 4e). However, significantly lower sub-genomic viral loads (Fig. 4f) and unmeasurable numbers of live infectious viruses (Fig. 4g) were achieved only by ZCB11. These results demonstrated that ZCB11 conferred significant protection against both Delta and Omicron variants, whereas ZB8 exhibited only partial protection against Delta but not Omicron. These findings were consistent with *in vitro* neutralizing activities of ZCB11 and ZB8 against live Delta and Omicron variants, respectively (Fig. 2c). Since the number of infectious viruses in the PBS group of Delta-challenged hamsters was over one order of magnitude higher than that in the PBS group of Omicron-challenged animals (Fig. 4d and 4g), higher amount of ZCB11 might be needed for improved suppression of sub-genomic viral loads against the Delta variant.

**Fig. 4.**
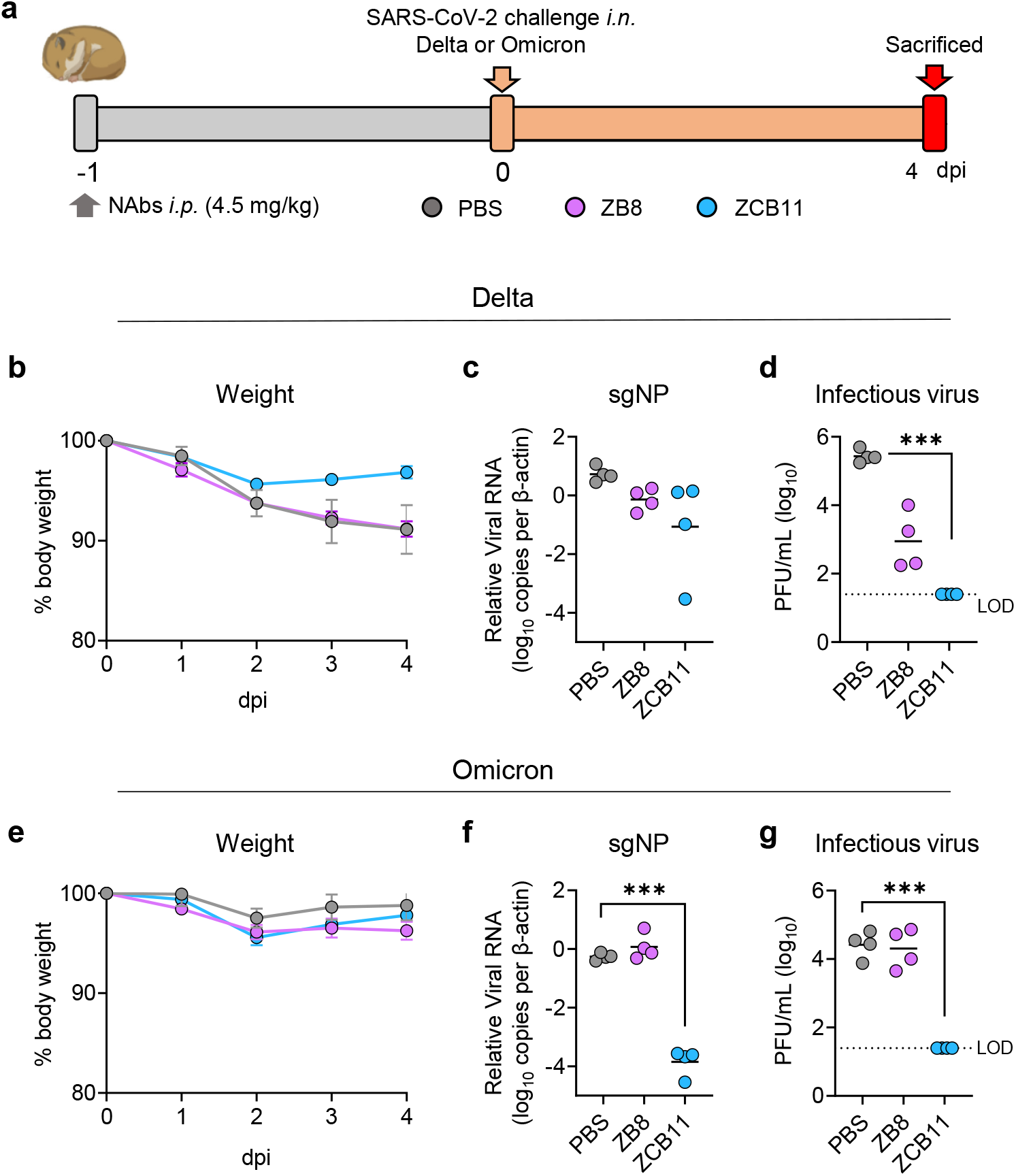
Efficacy of ZCB11 against authentic SARS-CoV-2 Delta and Omicron in golden Syrian hamsters as compared with ZB8. (**a**) Experimental schedule and color coding for different treatment groups. Three groups of hamsters (n=8) received a single intraperitoneal injection of PBS (grey), 4.5 mg/kg of ZB8 (purple) or 4.5 mg/kg of ZCB11 (blue) at one day before viral infection (−1 dpi). 24 hours later (day 0), each group was divided into two subgroups for intranasal challenge with 10^5^ PFU live SARS-CoV-2 Delta and Omicron variants, respectively. All animals were sacrificed on day 4 for final analysis. (**b, e**) Daily body weight was measured after viral infection. (**c, f**) The nucleocapsid protein (NP) subgenomic RNA copy numbers (normalized by β-actin) in lung homogenates were determined by a sensitive RT PCR. (**d, g**) Live viral plaque assay was used to quantify the number of infectious viruses in lung homogenates. Log10-transformed plaque-forming units (PFU) per mL were shown for each group. LOD: limit of detection. Each symbol represents an individual hamster with a line indicating the mean of each group. The color coding was consistently used in each graph. Statistics were generated using one-way ANOVA followed by Tukey’s multiple comparisons test. **p<0.01; ***p<0.001.

## DISCUSSION

It remains unclear what type of human monoclonal NAbs can potently neutralize all current SARS-CoV-2 VOCs including the Omicron Variant. In this study, we showed that the standard two-dose BNT162b2 vaccination was able to induce spike-specific memory B cells, from which we successfully cloned the elite bNAb ZCB11 around 130 days post the second vaccination. We demonstrated that ZCB11 not only neutralized all authentic SARS-CoV-2 VOCs including Omicron and Omicron_R346K_ at comparable high potency *in vitro* but also protected golden Syrian hamsters against the major circulating Omicron and Delta variants. Till now, few existing NAbs under clinical development have displayed similar neutralization breadth and *in vivo* potency ^20,21^. Since sequence analysis revealed that ZCB11 was a family member of public antibodies with the IGHV1-58 heavy chain and IGKV3-20 light chain, our findings have significant implication to vaccine design for inducing high amounts of ZCB11-like bNAb for broad protection and for clinical development of ZCB11-based immunotherapy against the pandemic SARS-CoV-2 VOCs.

ZCB11 overcomes naturally occurred spike mutations and deletions across current SARS-CoV-2 VOCs. Alpha variant with D614G and N501Y mutations enhanced RBD binding to human ACE2 receptor, transforming it into the most prevalent variant at the early stage of 2021 ^35^. N501Y alone was found conferring partial resistance to RBD-specific class I 910-30 and NTD-specific 4-18 NAbs ^21^. Subsequently, Beta, Gamma and Delta variants displayed the most troublesome mutations including K417N, E484K/Q/A and N501Y, conferring high resistance to RBD-specific class I and class II NAbs ^21,36-38^. E484K/Q/A led to almost complete loss of neutralization by potent RBD-specific class II LY-CoV555 and 2-15 ^21^. Attributed probably by antibody evasion, Delta variant, carrying L452R/T478K/D614G/P681R mutations, were found in more than 170 countries and accounted for 99% of newly confirmed cases before the Omicron variant ^39-41^. After the emergence of the Omicron variant with more than 30 mutations in viral spike protein ^42^, the ongoing wave of COVID-19 pandemic has already been dominated by it over the Delta variant in many countries probably due to further antibody evasion to almost all current vaccines and NAbs including those approved for clinical use or emergency use ^11,21,31,43^. N440K and G446S in Omicron conferred resistance to class III antibodies such as REGN10987 and 2-7 ^21^. G142D and del143-145 led to resistance to NTD-specific 4-18 and 5-7, whereas S371L conferred much broader resistance to RBD-specific class I, class III and class IV NAbs including potent Brii-196, REGN10987 and Brii-198 in clinical development ^21^. In this study, we consistently found that E484K/Q/A in Beta, Delta and Omicron variants conferred strong resistance to our RBD-specific class II ZB8 NAb. These resistant mutations, however, did not affect the potency of ZCB11 significantly. Although S371L in Omicron displayed partial resistance (∼11-fold) to ZCB11, similar amount of resistance was not observed against Omicron and Omicron_R346K_ that also contained S371L (Fig. 2c). Although ZCB11 shared 86.5% amino acid identity in variable regions with the previously reported ultrapotent S2E12 bNAb ^32,44^, it is critical to solve the real structure of RBD-ZCB11 Fab complex in future studies to understand if ZCB11 and S2E12 use an identical mode of action, which will be useful for novel vaccine design to elicit ZCB11-like bNAb responses.

Most public antibodies were RBD-specific class I NAbs ^21,29,30,45,46^. Accordingly, public antibody is encoded by B cell clonotypes isolated from different individuals that share similar genetic features ^47^. In a previous study, 7 of 13 NAbs were found using IGHV3-53/3-66 heavy chain and paired predominantly with IGKV1-9*01 light chain ^21^. These NAbs displayed abolished neutralizing activity after K417N in Beta variant was introduced into the pseudovirus neutralization assay. Interestingly, our newly cloned NAbs ZCB3, ZCC10 and ZCD3 utilized the same IGHV3-53/3-66 heavy chain but paired with IGKV1-9, IGKV3-20 and IGKV1-27 light chains, respectively (Supplementary Table 4). ZCB3, our second best bNAb, used the identical pair of IGHV3-53/3-66 and IGKV1-9 but did not display neutralization reduction against the K417N pseudovirus. ZCB3, however, showed reduced neutralization potency for over 10-fold against Beta and Omicron variants as compared with ZCB11. More interestingly, our elite bNAb ZCB11 used IGHV1-58 heavy chain and IGKV3-20 light chain, which also belongs to public antibodies reported by other groups ^29,32,47,48^. In these studies, patient-derived S2E12 and vaccine-induced 2C08 NAbs that shared 95% amino acid identity also used the same IGHV1-58 heavy chain and IGKV3-20 light chain. 2C08 was able to prevent challenges against Beta and Delta variants in the hamster model. Like 82.2% amino acid identity between ZCB11 and S2E12, ZCB11 and 2C08 shared 83.8% amino acid identity in their heavy chain variable regions. Their potency difference for neutralizing Omicron remained to be determined. Nevertheless, vaccine design in eliciting high amounts of ZCB11-like bNAb should be considered as a research priority, especially after its clonotype has been found in different ethnic human populations but have not been abundantly induced by current vaccines. Since ZCB11 protected hamsters against both the Delta and Omicron variants, the most dominant circulating SARS-CoV-2 VOCs in the world, our findings warrant the clinical development of ZCB11 and ZCB11-like bNAbs for patient immunotherapy and transmission prevention.

### Limitations of the study

ZCB11 probably represents the broadest breadth among bNAbs reported thus far with comparable potency against all current SARS-CoV-2 VOCs including Omicron and Omicron_R346K_. We are still in the process in determining its mode of action by solving structures of the RBD-ZCB11 Fab complex. Such information will be useful to guide vaccine design as mentioned because the frequency of elite vaccine remains low (2/34 in this study). To understand the frequency of ZCB11-like bNAb among BNT162b2-vaccinees, we need to investigate other elite responders who show equally potent bNAb responses. More ZCB11-like bNAbs should be also discovered to improve current antibody-based cocktail immunotherapy. For animal challenge experiments, we have done a single dose efficacy experiment. Different doses and routes of administration or antibody combination will be tested in future experiments to provide useful information to support clinical development of ZCB11 and ZCB11-like bNAb.

## METHODS

### Human subjects

A cohort of 34 vaccinees who received two doses of BNT162b2 before June 2021 were recruited for this study. The exclusion criteria include individuals with (1) documented SARS-CoV-2 infection, (2) high-risk infection history within 14 days before vaccination, (3) COVID-19 symptoms such as sore throat, fever, cough and shortness of breath. Clinical and laboratory findings were entered into a predesigned database. Written informed consent was obtained from all study subjects. This study was approved by the Institutional Review Board of The University of Hong Kong/Hospital Authority Hong Kong West Cluster (Ref No. UW 21-120 452).

### Viruses

Authentic SARS-CoV-2 D614G (MT835143), Alpha (MW856794), Beta (GISAID: EPI_ISL_2423556), Omicron (hCoV-19/Hong Kong/HKU-344/2021; GISAID accession number EPI_ISL_7357684) and Delta (hCoV-19/Hong Kong/HKU-210804-001/2021; GISAID: EPI_ISL_3221329) variants were isolated from respiratory tract specimens of laboratory-confirmed COVID-19 patients in Hong Kong ^24^. All experiments involving live SARS-CoV-2 followed the approved standard operating procedures of the Biosafety Level 3 facility at The University of Hong Kong ^49,50^.

### Cell lines

HEK293T cells, HEK293T-hACE2 cells and Vero-E6-TMPRSS2 cells were maintained in DMEM containing 10% FBS, 2 mM L-glutamine, 100 U/mL/mL penicillin and incubated at 37 □in a 5% CO2 setting ^51^. Expi293FTM cells were cultured in Expi293TM Expression Medium (Thermo Fisher Scientific) at 37 □in an incubator with 80% relative humidity and a 5% CO2 setting on an orbital shaker platform at 125 ±5 rpm/min (New Brunswick innova™ 2100) according to the manufacturer’s instructions.

### ELISA analysis of plasma and antibody binding to RBD and trimeric spike

The recombinant RBD and trimeric spike proteins derived from SARS-CoV-2 (Sino Biological) were diluted to final concentrations of 1 μg/mL/mL, then coated onto 96-well plates (Corning 3690) and incubated at 4 °C overnight. Plates were washed with PBS-T (PBS containing 0.05% Tween-20) and blocked with blocking buffer (PBS containing 5% skim milk or 1% BSA) at 37 °C for 1 h. Serially diluted plasma samples or isolated monoclonal antibodies were added to the plates and incubated at 37 °C for 1 h. Wells were then incubated with a secondary goat anti-human IgG labelled with horseradish peroxidase (HRP) (Invitrogen) or with a rabbit polyclonal anti-human IgA alpha-chain labelled with HRP (Abcam) and TMB substrate (SIGMA). Optical density (OD) at 450 nm was measured by a spectrophotometer. Serially diluted plasma from healthy individuals or previously published monoclonal antibodies against SARS-CoV-2 (B8) were used as negative controls.

### Isolation of SARS-CoV-2 spike-specific IgG+ single memory B cells by FACS

RBD-specific single B cells were sorted as previously described ^52^. In brief, PBMCs from infected individuals were collected and incubated with an antibody cocktail and a His-tagged RBD protein for identification of RBD-specific B cells. The cocktail consisted of the Zombie viability dye (Biolegend), CD19-Percp-Cy5.5, CD3-Pacific Blue, CD14-Pacific Blue, CD56-Pacific Blue, IgM-Pacific Blue, IgD-Pacific Blue, IgG-PE, CD27-PE-Cy7 (BD Biosciences) and the recombinant SARS-CoV-2 Spike-His described above. Two consecutive staining steps were conducted: the first one used an antibody and spike cocktail incubation of 30 min at 4 °C; the second staining involved staining with anti-His-APC and anti-His-FITC antibodies (Abcam) at 4 °C for 30 min to detect the His tag of the RBD. The stained cells were washed and resuspended in PBS containing 2% FBS before being strained through a 70-μm cell mesh filter (BD Biosciences). SARS-CoV-2 spike-specific single B cells were gated as CD19+CD27+CD3-CD14-CD56-IgM-IgD-IgG+Spike+ and sorted into 96-well PCR plates containing 10 μL of RNAase-inhibiting RT-PCR catch buffer (1M Tris-HCl pH 8.0, RNase inhibitor, DEPC-treated water). Plates were then snap-frozen on dry ice and stored at −80 °C until the reverse transcription reaction.

### Single B cell RT-PCR and antibody cloning

Single memory B cells isolated from PBMCs of infected patients were cloned as previously described ^53^. Briefly, one-step RT-PCR was performed on sorted single memory B cell with a gene specific primer mix, followed by nested PCR amplifications and sequencing using the heavy chain and light chain specific primers. Cloning PCR was then performed using heavy chain and light chain specific primers containing specific restriction enzyme cutting sites (heavy chain, 5′-AgeI/3′-SalI; kappa chain, 5′-AgeI/3′-BsiWI). The PCR products were purified and cloned into the backbone of antibody expression vectors containing the constant regions of human Igγ1. The constructed plasmids containing paired heavy and light chain expression cassettes were co-transfected into 293T cells (ATCC) grown in 6-well plates.

Antigen-specific ELISA and pseudovirus-based neutralization assays were used to analyze the binding capacity to SARS-CoV-2 spike and the neutralization capacity of transfected culture supernatants, respectively.

### Genetic analysis of the BCR repertoire

Heavy chain and light chain germline assignment, framework region annotation, determination of somatic hypermutation (SHM) levels (in nucleotides) and CDR loop lengths (in amino acids) were performed with the aid of the NCBI/IgBlast tool suite (https://www.ncbi.nlm.nih.gov/igblast/). Sequences were aligned using Clustal W in the BioEdit sequence analysis package (Version 7.2). Antibody clonotypes were defined as a set of sequences that share genetic V and J regions as well as an identical CDR3.

### Antibody production and purification

The paired antibody VH/VL chains were cloned into Igγ and Igk expression vectors using T4 ligase (NEB). Antibodies produced from cell culture supernatants were purified immediately by affinity chromatography using recombinant Protein G-Agarose (Thermo Fisher Scientific) according to the manufacturer’s instructions, to purify IgG. The purified antibodies were concentrated by an Amicon ultracentrifuge filter device (molecular weight cut-off 10 kDa; Millipore) to a volume of 0.2 mL in PBS (Life Technologies), and then stored at 4 °C or - 80 °C for further characterization.

### Pseudovirus-based neutralization assay

The neutralizing activity of NAbs was determined using a pseudotype-based neutralization assay as we previously described ^54^. Briefly, The pseudovirus was generated by co-transfection of 293T cells with pVax-1-S-COVID19 and pNL4-3Luc_Env_Vpr, carrying the optimized spike (S) gene (QHR63250) and a human immunodeficiency virus type 1 backbone, respectively ^54^. Viral supernatant was collected at 48 h post-transfection and frozen at -80 °C until use. The serially diluted monoclonal antibodies or sera were incubated with 200 TCID50 of pseudovirus at 37 °C for 1 hour. The antibody-virus mixtures were subsequently added to pre-seeded HEK 293T-ACE2 cells. 48 hours later, infected cells were lysed to measure luciferase activity using a commercial kit (Promega, Madison, WI). Half-maximal (IC50) or 90% (IC90) inhibitory concentrations of the evaluated antibody were determined by inhibitor vs. normalized response --4 Variable slope using GraphPad Prism 8 or later (GraphPad Software Inc.).

### Neutralization activity of monoclonal antibodies against authentic SARS-CoV-2

The SARS-CoV-2 focus reduction neutralization test (FRNT) was performed in a certified Biosafety level 3 laboratory. Neutralization assays against live SARS-CoV-2 were conducted using a clinical isolate previously obtained from a nasopharyngeal swab from an infected patient ^55^. The tested antibodies were serially diluted, mixed with 50 μL of SARS-CoV-2 (1×10^3^ focus forming unit/mL, FFU/mL) in 96-well plates, and incubated for 1 hour at 37°C. Mixtures were then transferred to 96-well plates pre-seeded with 1×10^4^/well Vero E6 cells and incubated at 37°C for 24 hours. The culture medium was then removed, and the plates were air-dried in a biosafety cabinet (BSC) for 20 mins. Cells were then fixed with a 4% paraformaldehyde solution for 30 min and air-dried in the BSC again. Cells were further permeabilized with 0.2% Triton X-100 and incubated with cross-reactive rabbit sera anti-SARS-CoV-2-N for 1 hour at RT before adding an Alexa Fluor 488 goat anti-rabbit IgG (H+L) cross-adsorbed secondary antibody (Life Technologies). The fluorescence density of SARS-CoV-2 infected cells were scanned using a Sapphire Biomolecular Imager (Azure Biosystems) and the neutralization effects were then quantified using Fiji software (NIH).

### Antibody binding kinetics and competition between antibodies measured by Surface Plamon Resonance (SPR)

The binding kinetics and affinity of recombinant monoclonal antibodies for the SARS-CoV-2 RBD protein (SinoBiological) were analyzed by SPR (Biacore T200, GE Healthcare). Specifically, the SARS-CoV-2 RBD protein was covalently immobilized to a CM5 sensor chip via amine groups in 10mM sodium acetate buffer (pH 5.0) for a final RU around 250. SPR assays were run at a flow rate of 10 uL/min in HEPES buffer. For conventional kinetic/dose-response, serial dilutions of monoclonal antibodies were injected across the spike protein surface for 180s, followed by a 900s dissociation phase using a multi-cycle method. Remaining analytes were removed in the surface regeneration step with the injection of 10 mM glycine-HCl (pH 1.5) for 60s at a flow rate of 30 μl/min. Kinetic analysis of each reference subtracted injection series was performed using the Biacore Insight Evaluation Software (GE Healthcare). All sensorgram series were fit to a 1:1 (Langmuir) binding model of interaction. Before evaluating the competition between antibodies, both the saturating binding concentrations of antibodies for the immobilized SARS-CoV-2 RBD protein were determined separately. In the competitive assay, antibodies at the saturating concentration were injected onto the chip with immobilized spike protein for 120s until binding steady-state was reached. The other antibody also used at the saturating concentration was then injected for 120s, followed by another 120s of injection of antibody to ensure a saturation of the binding reaction against the immobilized RBD protein. The differences in response units between antibody injection alone and prior antibody incubation reflect the antibodies’ competitive ability by binding to the RBD protein.

### Model building of ZCB11 and structure presentation

A model of ZCB11 variable regions was generated based on the protein sequence by the SWISS-MODEL using the crystal structure of S2E12 Fab fragment (Research Collaboratory for Structural Bioinformatics [RCSB] PDB code 7K3Q) as the template. The structure alignment, cartoon representations, labeling of amino acids in RBD (from PDB 7K45) were generated by PyMOL.

### Hamster experiments

*In vivo* evaluation of monoclonal antibody ZB8 or ZCB11 in the established golden Syrian hamster model of SARS-CoV-2 infection was performed as described previously, with slight modifications ^33^. The animal experiments were approved by the Committee on the Use of Live Animals in Teaching and Research (CULATR 5359-20) of the University of Hong Kong (HKU). Briefly, 6-10-week-old male and female hamsters were obtained from the Chinese University of Hong Kong Laboratory Animal Service Centre through the HKU Centre for Comparative Medicine Research. The hamsters were housed with access to standard pellet feed and water ad libitum until live virus challenge in the BSL-3 animal facility at Department of Microbiology, HKU. The viral challenge experiments were then conducted in our Biosafety Level-3 animal facility following SOPs strictly, with strict adherence to SOPs. The hamsters were randomized from different litters into experimental groups. Experiments were performed in compliance with the relevant ethical regulations ^33^. For prophylaxis studies, 24 hours before live virus challenge, three groups of hamsters were intraperitoneally administered with one dose of test antibody in phosphate-buffered saline (PBS) at the indicated dose. At day 0, each hamster was intranasally inoculated with a challenge dose of 100 μL of Dulbecco’s Modified Eagle Medium containing 10^5^ PFU of SARS-CoV-2 Delta variant or Omicron variant under anesthesia with intraperitoneal ketamine (200 mg/kg) and xylazine (10 mg/kg). The hamsters were monitored daily for clinical signs of disease. Syrian hamsters typically clear virus within one week after SARS-CoV-2 infection. Accordingly, animals were sacrificed for analysis at day 4 after virus challenge with high viral loads ^33^. Half the nasal turbinate, trachea, and lung tissues were used for viral load determination by quantitative RT-qPCR assay ^56^ and infectious virus titration by plaque assay ^33^ as we described previously.

### Quantification and statistical analysis

Statistical analysis was performed using PRISM 8.0 or later. Ordinary one-way ANOVA and multiple comparisons were used to compare group means and differences between multiple groups. Unpaired Student’s t tests were used to compare group means between two groups only. A P-value <0.05 was considered significant. The number of independent experiments performed, the number of animals in each group, and the specific details of statistical tests are reported in the figure legends and the Methods section.

## Supporting information

Supplemental meterials

## SUPPLEMENTAL INFORMATION

The supplemental information includes 7 Tables and 3 Figures.

## ACKNOWLEDGMENTS

This study was supported by the Hong Kong Research Grants Council Collaborative Research Fund (C7156-20G, C1134-20G and C5110-20G) and Shenzhen Science and Technology Program (JSGG20200225151410198 and JCYJ20210324131610027); the Hong Kong Health@InnoHK, Innovation and Technology Commission; and the China National Program on Key Research Project (2020YFC0860600, 2020YFA0707500 and 2020YFA0707504); and donations from the Friends of Hope Education Fund. Z.C.’s team was also partly supported by the Hong Kong Theme-Based Research Scheme (T11-706/18-N). This study was also partly supported by funding the Health and Medical Research Fund, the Food and Health Bureau, The Government of the Hong Kong Special Administrative Region (Ref no.: COVID1903010-Project 7 and 20190572); the Consultancy Service for Enhancing Laboratory Surveillance of Emerging Infectious Diseases and Research Capability on Antimicrobial Resistance for Department of Health of the Hong Kong Special Administrative Region Government; the National Program on Key Research Project of China (grant no. 2020YFA0707500 and 2020YFA0707504); Sanming Project of Medicine in Shenzhen, China (grant no. SZSM201911014); the High Level-Hospital Program, Health Commission of Guangdong Province, China; the Major Science and Technology Program of Hainan Province (ZDKJ202003); and the research project of Hainan academician innovation platform (YSPTZX202004); and donations from the Shaw Foundation of Hong Kong, the Richard Yu and Carol Yu, Michael Seak-Kan Tong, May Tam Mak Mei Yin, Lee Wan Keung Charity Foundation Limited, the Providence Foundation Limited (in memory of the late Lui Hac Minh), Hong Kong Sanatorium & Hospital, Hui Ming, Hui Hoy and Chow Sin Lan Charity Fund Limited, Chan Yin Chuen Memorial Charitable Foundation, Marina Man-Wai Lee, the Hong Kong Hainan Commercial Association South China Microbiology Research Fund, the Jessie & George Ho Charitable Foundation, Perfect Shape Medical Limited, Kai Chong Tong, Tse Kam Ming Laurence, Foo Oi Foundation Limited, Betty Hing-Chu Lee, Ping Cham So, and Lo Ying Shek Chi Wai Foundation. The funding sources had no role in the study design, data collection, analysis, interpretation, or writing of the report. Finally, we thank Dr. David D. Ho for kindly providing the expression plasmids encoding for D614G, B.1.1.7 and B.1.351 variants and Dr. Linqi Zhang for B.1.617.2.

## AUTHOR CONTRIBUTIONS

Conceptualization, Z.C.; HuNAb cloning, Z.B.; experimental design, Z.C., Z.B., R.Z., J.F.-W.C.; hamster experiments, J.F.-W.C., V.K.-M.P., C.C.-S.C., J.O.-L.T.,, C.C.-Y.C.; clinical specimens, Q.P., K.K.-W.T.; pseudovirus neutralization assay, M.L., Q.P., B.C.; authentic virus neutralizing assay, R.Z., B.W.-Y.M., P.W., H.C., L.L., H.C.; viral plaque assay, S.Y.; plasmid cloning, M.L., B.C., H.-O.M. viral RNA measurement, K.-K.A.; critical comments, supports and materials, K.-Y.Y.

## DECLARATION OF INTEREST

J.F.-W.C. has received travel grants from Pfizer Corporation Hong Kong and Astellas Pharma Hong Kong Corporation Limited and was an invited speaker for Gilead Sciences Hong Kong Limited and Luminex Corporation. The funding sources had no role in study design, data collection, analysis or interpretation or writing of the report. The other authors declare no conflicts of interest except for a provisional patent application filed for human monoclonal antibodies generated in our laboratory. Z.C., B.Z., and R.Z. are the co-inventers of NAbs ZCB11, ZCB3 and ZCD3.

### Reporting Summary

Further information on research design is available in the Nature Research Reporting Summary linked to this article.

### Data availability

The data of this studies are available upon reasonable request and accession codes will be available before publication.

### Code availability

No custom computer code or algorithm used to generate results that are reported in the paper and central to its main claims.

