## Supplemental meterials for "An elite broadly neutralizing antibody protects SARS-CoV-2 Omicron variant challenge": Supplementary figures and tables 20210105.pdf

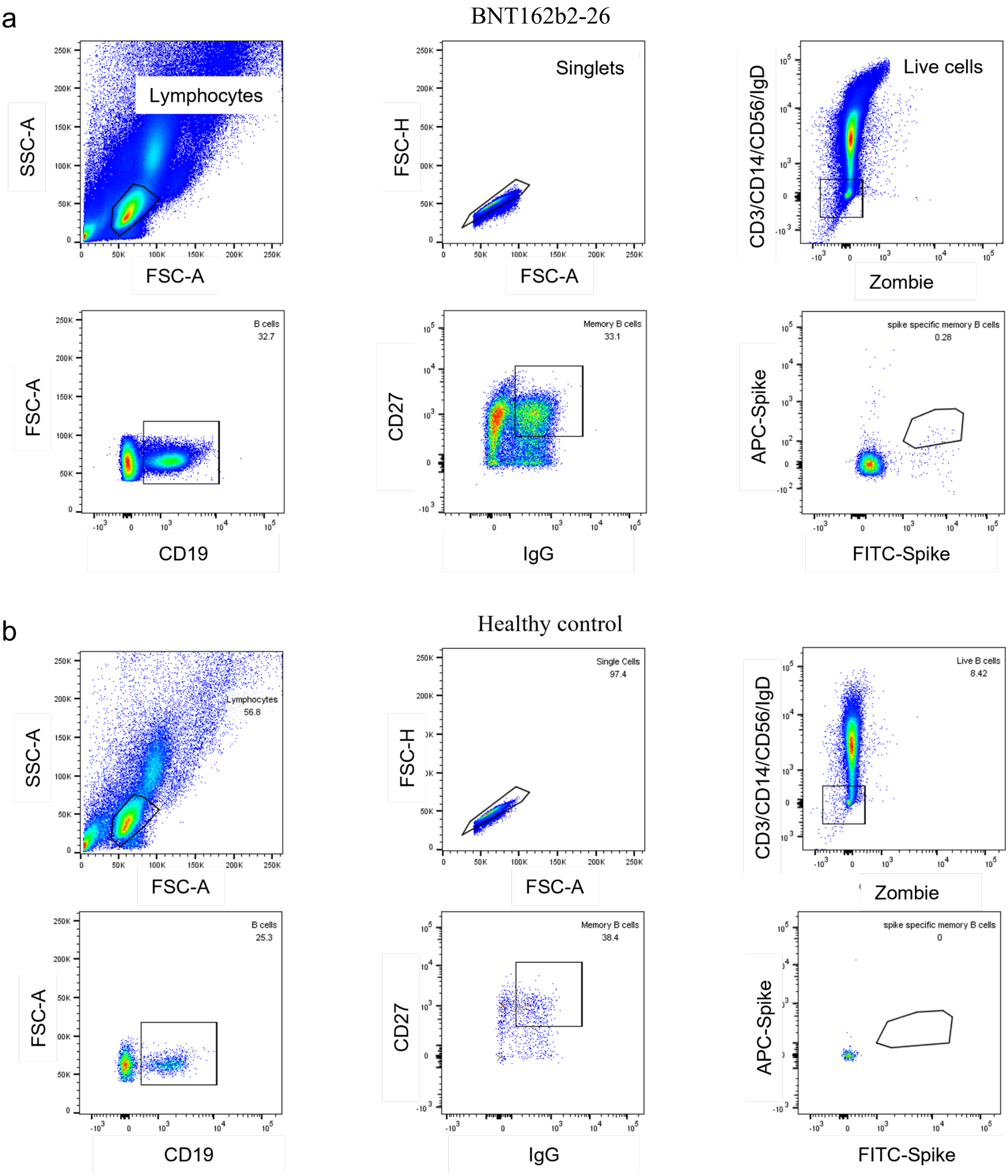


**Supplementary Fig. 1. Gating strategy for sorting antigen specific memory B cells from the BNT162b2-26 vaccinee (a) as compared with a healthy control (b).**

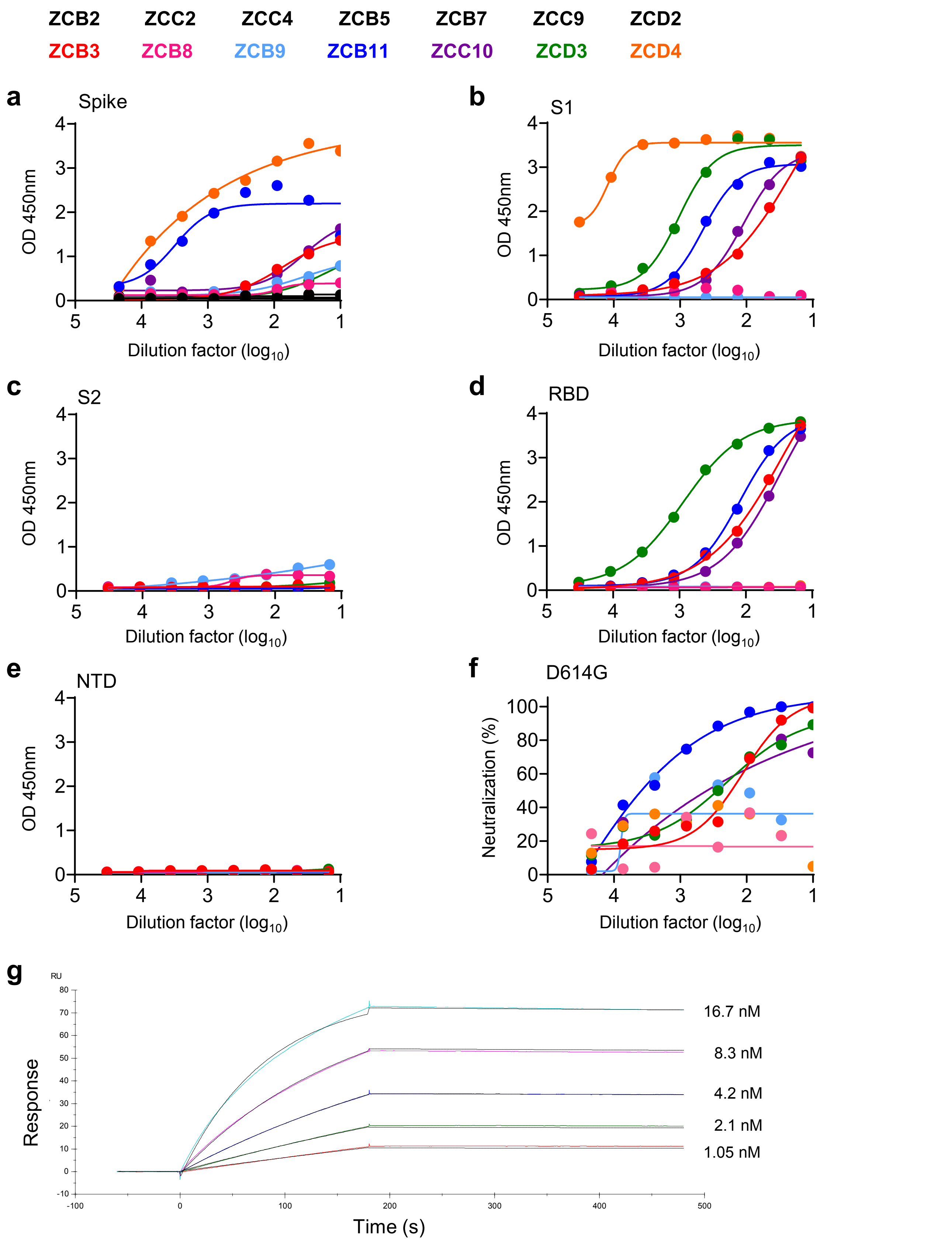


**Supplementary Fig 2. Binding and neutralizing activities of 14 newly cloned human monoclonal antibodies. (a-f)** HEK 293T cells were transfected with expression plasmids encoding paired heavy and light chains. Two days after transfection, culture supernatants were subjected to binding test to SARS-CoV-2 Spike (**a**), S1 (**b**), S2 (**c**), RBD (**d**) and NTD (**e**) by ELISA, respectively. (**f**) Neutralization activities of culture supernatants were also determined by the pseudotyped SARS-CoV-2 WT in 293T-ACE2 cells. (**g**) Antibody binding kinetics by SPR. ZCB11 was captured on protein A covalently immobilized onto a CM5 sensor chip followed by injection of purified soluble SARS-CoV-2 WT RBD at five different concentrations. The black lines indicated the experimentally derived curves while the color lines represented fitted curves based on the experimental data.

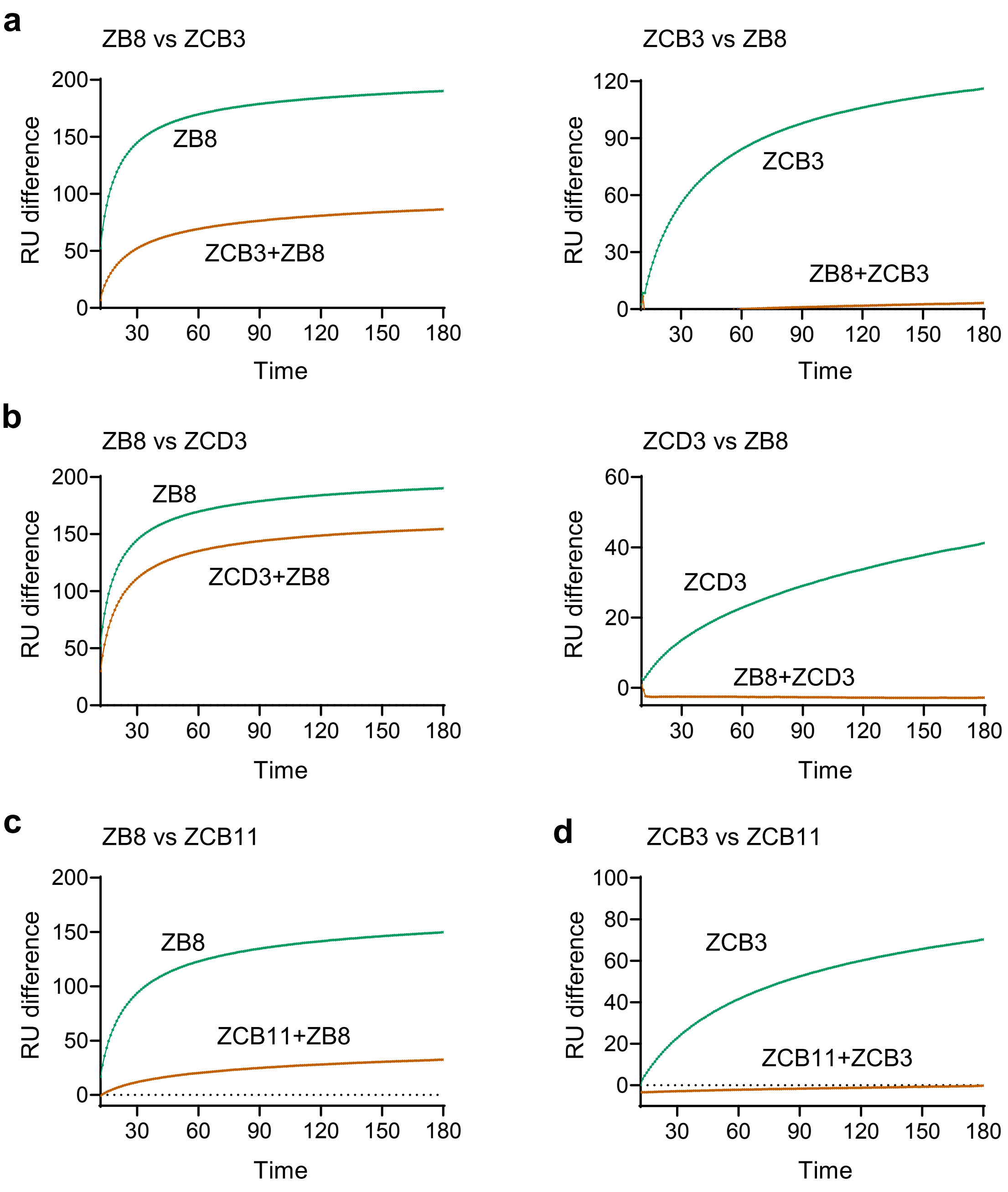


**Supplementary Fig 3. Competition binding assay of newly cloned NAbs with ZB8**. (**a-d**) The sensorgrams show distinct binding patterns when pairs of testing antibodies were sequentially applied to the purified SARS-CoV-2 RBD covalently immobilized onto a CM5 sensor chip. Color coding curves indicate distinct binding patterns of representative NAbs to RBD with (orange) or without (green) prior incubation with each testing antibody.

Supplementary Table 1. Characteristics of BNT162b2 vaccinees

| Vaccinee ID | Age | Gender | Vaccination doses | Sample collection (days post 2nd vaccination) |
|---|---|---|---|---|
| BioNTech-1 | 28 | F | 2 | 30 |
| BioNTech-2 | 29 | M | 2 | 31 |
| BioNTech-3 | 31 | M | 2 | 31 |
| BioNTech-4 | 22 | M | 2 | 30 |
| BioNTech-5 | 36 | M | 2 | 31 |
| BioNTech-6 | 32 | M | 2 | 31 |
| BioNTech-11 | 26 | M | 2 | 24 |
| BioNTech-12 | 27 | F | 2 | 31 |
| BioNTech-13 | 35 | F | 2 | 24 |
| BioNTech-16 | 25 | F | 2 | 31 |
| BioNTech-18 | 29 | F | 2 | 31 |
| BioNTech-19 | 30 | M | 2 | 30 |
| BioNTech-20 | 27 | F | 2 | 30 |
| BioNTech-21 | 52 | F | 2 | 31 |
| BioNTech-24 | 35 | F | 2 | 22 |
| BioNTech-25 | 35 | M | 2 | 22 |
| BioNTech-26 | 58 | M | 2 | 7 |
| BioNTech-27 | 29 | F | 2 | 26 |
| BioNTech-28 | 25 | M | 2 | 26 |
| BioNTech-35 | 25 | F | 2 | 14 |
| BioNTech-37 | 33 | F | 2 | 47 |
| BioNTech-38 | 20 | M | 2 | 47 |
| BioNTech-39 | 32 | F | 2 | 25 |
| BioNTech-42 | 32 | M | 2 | 47 |
| BioNTech-46 | 25 | M | 2 | 42 |
| BioNTech-55 | 25 | M | 2 | 12 |
| BioNTech-57 | 52 | M | 2 | 18 |
| BioNTech-58 | 47 | F | 2 | 11 |
| BioNTech-59 | 44 | F | 2 | 17 |
| BioNTech-62 | 30 | F | 2 | 37 |
| BioNTech-63 | 29 | F | 2 | 47 |
| BioNTech-64 | 66 | F | 2 | 36 |
| BioNTech-67 | 50 | F | 2 | 12 |
| BioNTech-68 | 33 | M | 2 | 43 |

Supplementary Table 2. Neutralization titers of BNT162b2-26 plasma

| Sample | Neutralization titers ($IC_{50}$) | | | | | |
|---|---|---|---|---|---|---|
| | WT | Alpha | Beta | Gamma | Delta | Omicron |
| BNT162b2-26 | 899 | 982 | 5085 | 466 | 2229 | 115 |
| Average | 731 | 474 | 95 | 532 | 164 | 35 |

Supplementary Table 3. Epitopes of four isolated SARS-CoV-2 specific antibodies

| Antibody | Epitope |
|---|---|
| ZCB3 | RBD |
| ZCB8 | S2 |
| ZCB9 | S2 |
| ZCB11 | RBD |
| ZCC10 | RBD |
| ZCD3 | RBD |
| ZCD4 | S1 |

Supplementary Table 4. Gene family analysis of four neutralizing antibodies

| NAbs | Heavy chain | | | | Light chain | | | |
|---|---|---|---|---|---|---|---|---|
| | IGHV | IGHJ | CDR3 length | SHM (%) | IGKV | IGKJ | CDR3 length | SHM (%) |
| ZCB3 | IGHV3-53*04 | IGHJ6*02 | 12 | 5.1 | IGKV1-9*01 | IGKJ2*01 | 9 | 3.2 |
| ZCB11 | IGHV1-58*02 | IGHJ3*02 | 16 | 5.5 | IGKV3-20*01 | IGKJ1*01 | 9 | 2.8 |
| ZCC10 | IGHV3-53*04 | IGHJ6*02 | 12 | 5.1 | IGKV3-20*01 | IGKJ4*01 | 8 | 1.7 |
| ZCD3 | IGHV3-66*01 | IGHJ4*02 | 16 | 3.8 | IGKV1-27*01 | IGKJ1*01 | 10 | 1.4 |

Supplementary Table 5. Binding ability of public NAbs to SARS-CoV-2 RBD and spike.

| NAbs | SARS-CoV-2 RBD | SARS-CoV-2 spike |
|---|---|---|
| | $EC_{50}$ (μg/mL) | $EC_{50}$ (μg/mL) |
| ZCB3 | 0.027 | 0.092 |
| ZCB11 | 0.020 | 0.020 |
| ZCC10 | 0.041 | 0.245 |
| ZCD3 | 0.156 | 1.582 |

Supplementary Table 6. Surface plasmon resonance analysis of ZCB11

| Curve | Conc (M) | ka (1/Ms) | kd (1/s) | KD (M) | Rmax (RU) | tc |
|---|---|---|---|---|---|---|
| 0.15625 μg/mL | 1.04E-09 | | | | | |
| 0.3125 μg/mL | 2.08E-09 | | | | | |
| 0.625 μg/mL | 4.17E-09 | | | | | |
| 1.25 μg/mL | 8.33E-09 | | | | | |
| 2.5 μg/mL | 1.67E-08 | | | | | |
| | | 7.35E+05 | 4.22E-05 | 5.75E-11 | 81.21 | 9.11E+14 |

Supplementary Table 7. Neutralization $IC_{50}$ values of public NAbs.

| NAbs | Pseudovirus $IC_{50}$ (ng/mL) | | | | | | Live virus $IC_{50}$ (ng/mL) | | | | | |
|---|---|---|---|---|---|---|---|---|---|---|---|---|
| | WT | α | β | γ | Δ | ο | WT | α | β | γ | Δ | ο |
| ZCB3 | 40.7 | 16.1 | 57.7 | 37.8 | 77.9 | 531.6 | 176.3 | 312.5 | 1383 | 540.6 | 41.3 | 6450 |
| ZCB11 | 5.2 | 8.9 | 6.1 | 34.5 | 31.5 | 6 | 51 | 85.1 | 39.9 | 56.9 | 11.2 | 36.8 |
| ZCC10 | 316.2 | 60.5 | *N.A.* | 1353 | 369.7 | *N.A.* | / | / | / | / | / | / |
| ZCD3 | 355.8 | 210.1 | *N.A.* | 344.7 | 141.2 | *N.A.* | 2358 | *N.A.* | *N.A.* | *N.A.* | 392.6 | *N.A.* |

*N.A.*: Not applicable
